# ASD: Antigen-Specific Antibody Database

**DOI:** 10.1101/2025.11.24.690097

**Authors:** Arkadiusz Czerwiński, Paweł Dudzic, Konrad Wójtowicz, Igor Jaszczyszyn, Weronika Bielska, Sonia Wrobel, Samuel Demharter, Roberto Spreafico, Victor Greiff, Konrad Krawczyk

## Abstract

The development of computational models addressing therapeutic antibodies faces significant challenges due to the scarcity of data. A critical data element is the set of antibody-antigen interaction pairs associated with sequences. To address this issue, we developed the Antigen Specific Antibody Database (ASD, https://naturalantibody.com/agab/), a database aggregating antibody-antigen interaction data from multiple studies with standardized formatting and annotations. Our dataset compilation strategy resulted in data from 15 distinct sources, resulting in 1,097,946 unique antibody-antigen interactions (with 9,575 unique antigens). The ASD database captures diverse affinity measures and qualitative binding assessment, along with metadata including UniProt and PDB identifiers, target protein names, confidence levels, and experimental conditions. Through this integration drive, we make available an ample resource of interaction data gathered from the public domain to act as a foundation for model development and further data generation.

## Introduction

Antibodies are essential components of the immune system, capable of recognizing and binding specific molecular targets on pathogens with high precision ^1,2^. Their ability to distinguish a vast range of antigens is driven by sequence variability in the Complementarity Determining Regions (CDRs) directly interacting with targets. This specificity enables antibodies to block harmful activity or mark threats for immune clearance, making them powerful tools in natural immunity and therapeutic design ^3^.

In recent decades, therapeutic antibodies have emerged as a cornerstone of modern biomedicine ^4^. The development of monoclonal antibody technology has enabled the generation of highly specific agents that can be directed against disease-associated antigens, leading to targeted treatments with fewer off-target effects ^5^. The process typically begins with the identification of relevant targets, followed by the selection of candidate antibodies with desired binding properties. These candidates are subsequently engineered to optimize stability, binding affinity, and pharmacokinetics while reducing immunogenicity ^6^. Central to these engineering efforts are modifications to the antibody variable regions, with a focus on refining the structural and chemical properties of the CDRs ^7^.

Despite significant technological advances, the success rate in antibody engineering and biological drug development remains modest ^8,9^. While many antibody candidates demonstrate strong potential in vitro, only a fraction ultimately progress through preclinical and clinical stages to achieve regulatory approval ^10^. Industry-wide analyses based on comprehensive data show that the overall likelihood of a biologic drug progressing from Phase I clinical trials to market approval is approximately 14.3%, with success rates among leading pharmaceutical companies ranging broadly from 8% to 23% ^11^. These figures highlight the complexity and high attrition rates characteristic of the field. However, recent breakthroughs, such as deep learning-based structure prediction, high-throughput screening technologies, and synthetic biology approaches are streamlining the design and refinement of therapeutic antibodies ^12–14^. The innovations are poised to improve efficiency, shorten development timelines, and potentially increase the success rate of antibody-based therapies in the coming years.

A major obstacle to fully realizing the potential of these computational methods is the scarcity of available antibody-antigen interaction data ^15–17^. A number of purpose-built affinity datasets revealed the limitations of binding prediction methods ^17–19^, suggesting efforts in data curation/generation rather than model development. Currently, binding data are dispersed across disparate sources, generated using varying experimental protocols, and reported in inconsistent formats. This heterogeneity hampers large-scale analyses and limits the ability to train generalizable machine learning models capable of accurate prediction across diverse antibody-antigen pairs. The previous efforts at consolidating such data in a domain-specific manner include databases such as the Structural Antibody Database (SAbDab) ^20^, PLAbDab ^21^, which identify antibody data in the PDB and GenBank, respectively. Although much work has been put into creating these resources, the data therein still requires additional preprocessing to obtain a clean dataset of binder pairs.

Here, we developed the Antigen-Specific Antibody Database (ASD) as a unified repository of antibody–antigen interactions. We strived to provide researchers with a self-sufficient dataset that would not require excessive preprocessing by aggregating suitable databases and enriching them with proper metadata, along with confidence levels that reflect the quality of sequences of provided bindings, along with an empirical assessment of binding. In addition to confidence levels, we provide additional lineage metadata in the form of target UniProt and/or PDB IDs relating to the target. In case of patent/literature databases, it points to the specific patents/publications where possible, and in case of aggregate databases, they point to the original paper the data comes from. In addition, each major dataset is described in the paper along with citations and links, which enables further filtering and reproducibility. Not all datasets present in this study share this same type of affinity measurements, resulting in a diverse, heterogeneous dataset that matches an array of use cases without the explicit need to search for external resources.

Altogether, by aggregating data from multiple sources and applying consistent formatting and annotation standards, ASD aims to provide a comprehensive, accessible, and high-quality resource of antibody-binders from the public domain.

## Materials and Methods

### Dataset aggregation

To construct a large and integrated dataset of antibody-antigen interactions, we implemented a diversified data collection strategy that combined information from open-access databases, peer-reviewed publications, and previously curated datasets cited in the literature. For each selected source, we extracted fundamental biological and biophysical characteristics of antibody-antigen interactions, focusing particularly on obtaining complete amino acid sequences of antibody heavy and light chains. We could not extend this condition to the sequence of the antigens that are oftentimes referred to only by their names.

It comes as no surprise that ASD partially overlaps with several established antibody-antigen resources. For instance, the Antibinder dataset refines and expands upon earlier datasets, so its integration includes older versions of some datasets that we can obtain directly as well. Following the filtering and quality-control procedures applied during Antibinder’s construction, approximately 50% of the original entries were retained, as seen in the overlap between ASD, Antibinder, and their respective source datasets. Specifically, ASD shares 2,018 PDB targets and 459 of 10,540 unique antibodies with SAbDab, and 2,977 antibodies with PLAbDab. Dataset intersections were determined through direct sequence and structure comparisons without prior alignment, which may have slightly reduced matching ratios. In Table 1, we summarized the differences of ASD with respect to some leading antibody datasets.

**Table 1:** Comparison between leading datasets in antibody sequence engineering and ASD based on their features and completeness.

| Dataset name | Source completeness | Target completeness | Preprocessing required |
| --- | --- | --- | --- |
| CovAbdab<sup>22</sup> | Complete Vh or VHH sequence | Only target name | No preprocessing required |
| PLabDab<sup>21</sup> | Complete sequences | Sometimes mentioned by name, with several targets lacking | Manual or automated search required for respective target sequence |
| SabDab<sup>20</sup> | Pdb structure, specific chains | Pdb structure, specific chain | Loading of the pdb/fastA required |
| ASD | Complete sequences | Complete sequence with additional metadata | No preprocessing required |

The final dataset was compiled from 15 distinct sources, resulting in 25 individual datasets (Table 2) representing a diverse array of antibody-antigen interaction records. The curation process involved a multi-stage pipeline combining automated extraction, normalization, and manual verification steps. Key data sources included structural databases (e.g., PDB), sequence-based repositories (e.g., GenBank), and literature-derived or patent datasets processed using natural language processing and optical character recognition (OCR) technologies.

**Table 2:**
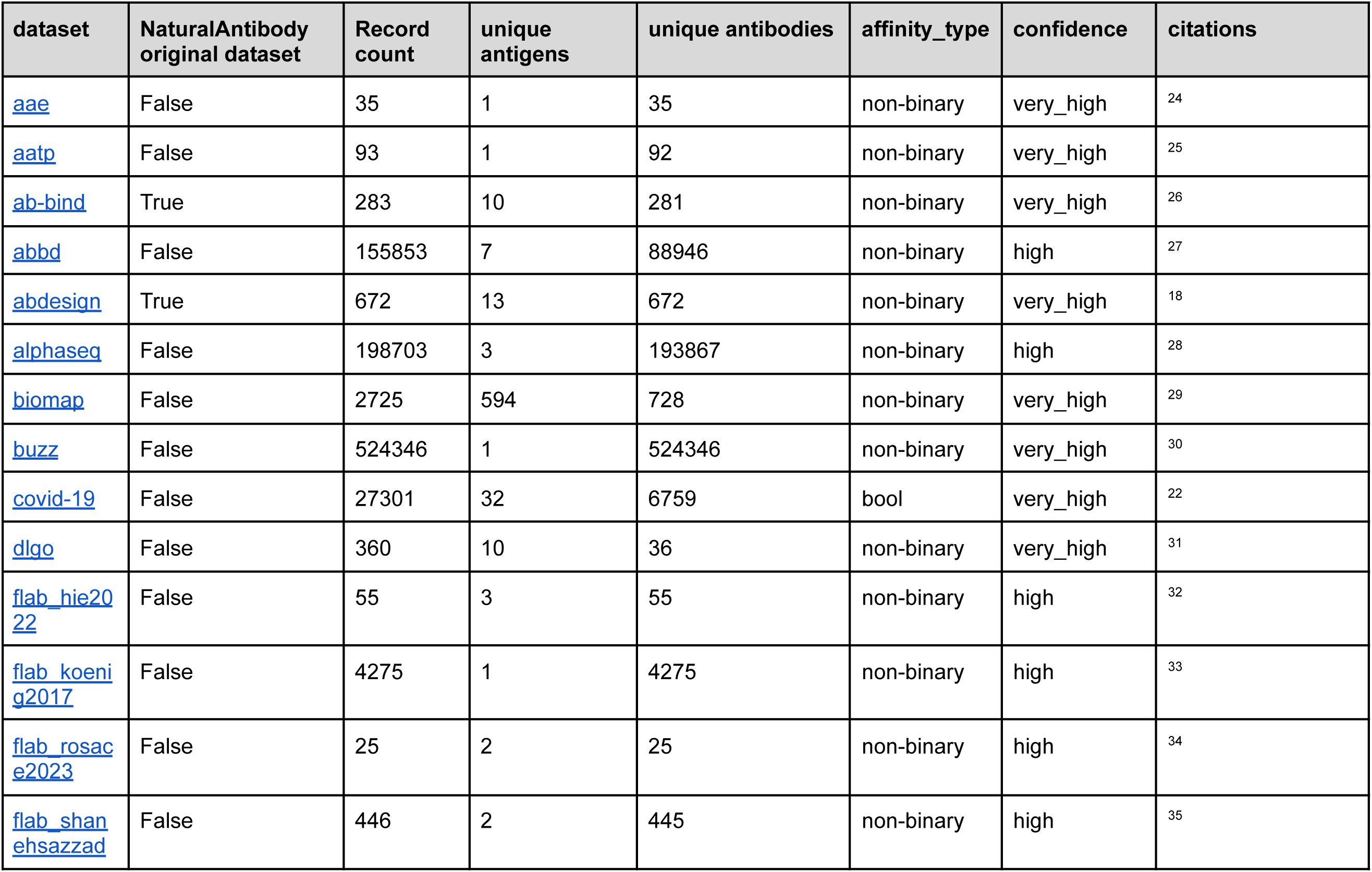
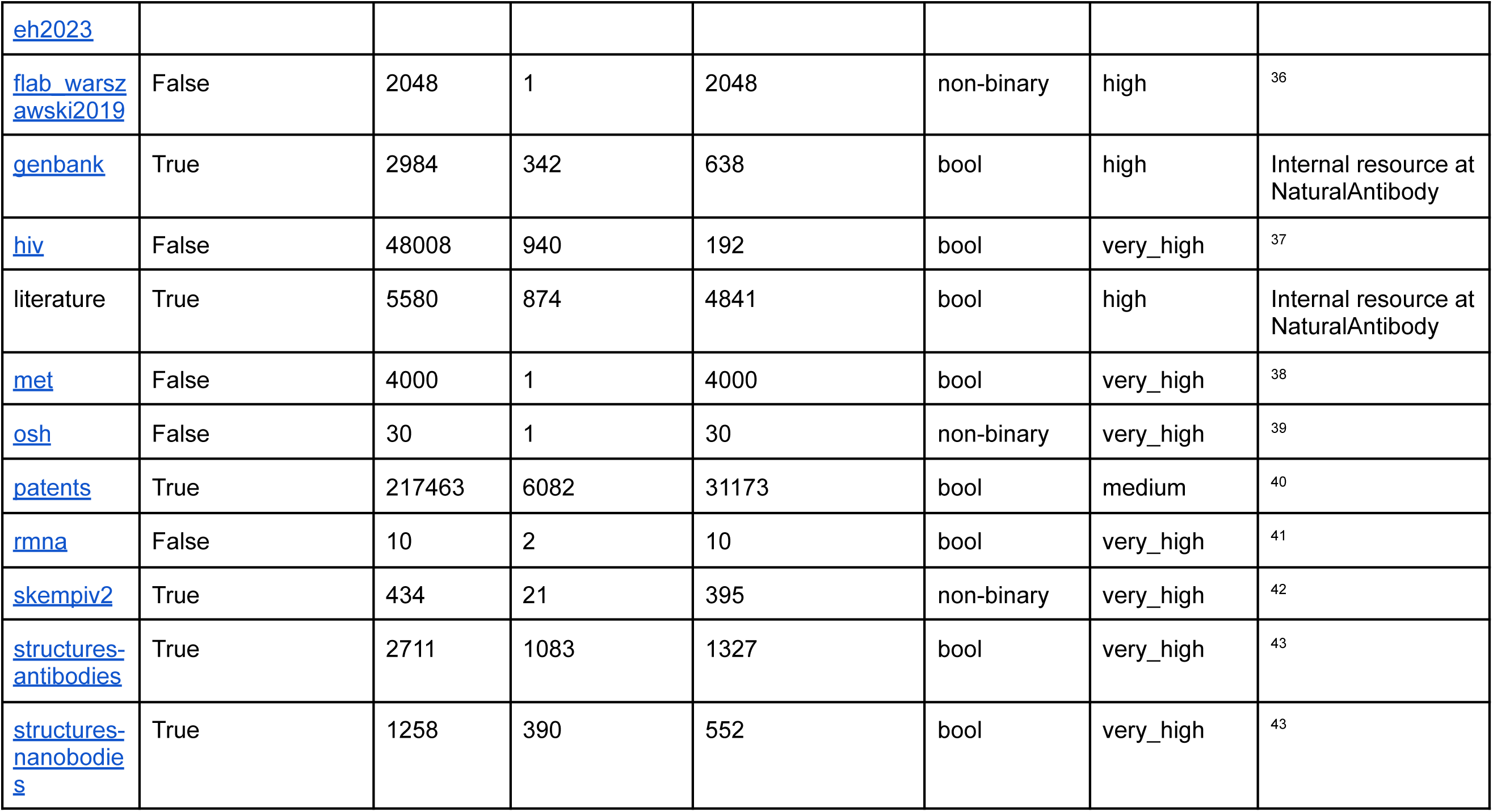
List of datasets used in the study. The datasets were categorized based on the level of confidence we have in the data. For instance, structural data is very high quality as the sequences are known down to the atomic level, whereas patent data is lower quality as one cannot be certain that the molecules were actually tested ^23^.

Some of the datasets did not directly include antigen or antibody sequences. Instead, the sequence data was present either in the form of a UniProt/PDB identifier or a respective protein name, possibly including additional mutation points. If the sequence was stored in the form of a UniProt/PDB ID, the full sequence was retrieved from a respective database, and its corresponding ID was added to the metadata. In many cases, the sequences had to be manually curated by applying mutations specified in the publication to the parental sequence.

The papers and samples that could not be reconstructed or the information regarding their binding was uncertain were removed from the dataset, leaving only curated and identifiable samples. Although most of the sources obtained from research papers were structured in their nature (e.g., available in tabular format), some of them required additional manual curation or application of Optical Character Recognition (OCR) techniques performed by Claude Opus 4.1 through a web interface.

### Dataset Curation

Each of the major sources was individually processed in a way that preserves the heterogeneity in input format, such as “affinity_type” or general target metadata. This ensured that each dataset could be analyzed separately, as well as being included in any permutation of aggregations with different datasets.

The **AAE** dataset, first introduced by Lena Erlach et al. ^24^ had its data parsed from supplementary tables in PDF format. After parsing, the outcome was manually verified, resulting in two sub-sources - sequence and affinity, which were then joined, forming the core of the dataset. Then, the target was assigned based on the name found in the paper. The affinity was expressed as a KD measurement [nM]. Similarly, the **AAPT** dataset underwent similar antibody processing, with the exception of performing OCR on the target and antibody sequences that have been placed in the image in the main paper. The antibody sequence for each sample was then obtained through respective mutations, which were then assigned an appropriate KD affinity. The next three datasets (**skempiv2**, **ab-bind**, **abdesign**) originated from the same already processed dataset - Naturalantibody’s AbDesign Database, with little to no preprocessing needed, with each affinity type already present in the database. The exact details on those datasets are present under a link: https://naturalantibody.com/ab-design/. The **ABBD** originated from a hugging face resource that aggregated affinities (measured originally as - log KD) for several PDB’s. In this particular case, the target required obtaining it through the scraping of associated fasta files in the RCSB PDB database. Aside from the target sequence, the affinity and antibody sequences were present in the original database. The **Alphaseq** dataset, originating from the Alphabind paper, presents the “alphaseq” affinity measurement method, which produced confidence in antibody-antigen binding. The dataset consists of two tables, one with affinity data and the other with associated target sequences. By combining those two tables and information about their classification (scFv and nanobody), the final dataset was condensed into a single table of unified format. **AntiBinder** dataset stems from a repository associated with the source paper. The dataset contained four already preprocessed datasets (**hiv**, **covid-19**, **met**, **biomap**), which were previously aggregated and filtered from other databases such as CovAbdab based on the presence of the target and affinity. The dataset gathered both boolean and numerical affinity measurements (expressed as delta_g). The tables were unified into common naming conventions, linked to the specific affinity type, and enriched with basic metadata. The biggest dataset in our work - **buzz** stemming from the paper “Baselining the buzz” is made by reading preprocessed CSV files, which were unified. After the unification, the full heavy sequence was obtained, while the rest of the sequences were obtained by the identification of corresponding sequences based on the paper. Similarly, the binding confidence was expressed as one of the “high” “medium” or “low” categories. The **DLGO** dataset contained all of its data from a single table involving affinity data and mutations. The table was unified as the CSV file via the usage of the Claude Opus model, which was used to prepare the correct structure and initial values. The mistakes in the OCR were then manually corrected and saved into a final CSV, which was then used to obtain full sequences from mutation points. The resulting dataset measured the binding affinity in ic_50 [μg/ml] format. The **Flab** dataset consists of five sub-datasets - **koenig2017**, **warszawski2019**, **flab_shanehsazzadeh2023**, **flab_hie2022**, **flab_rosace2023.** Datasets, although containing complete sequences and KD affinity data, lacked target sequences. This situation was remedied by assigning target sequences by names that appeared in the original papers, with a degree of confidence related to the paper. The **OSH** dataset is another resource which requires the use of OCR techniques, due to the entire table containing sequence and KD affinity appearing in image format. The OCR results were manually verified and parsed into CSV format. The data contained in **RMNA** required only slight schema adjustments, as it was perfectly contained in an Excel spreadsheet, with only boolean binding information.

**GenBank**, **Literature, Patents,** and **Structures** ^43^ databases were all obtained from internal NaturalAntibody resources. The datasets also include the same type of affinity - boolean - as the datasets contained only positive, binding examples. The **GenBank** dataset contains both nanobodies and monoclonal antibodies; they were tagged based on internal metadata and normalized to correct structure. In the case of the **Literature** database, it required fetching target sequences, which were obtained from either Uniprot, Uniparc, or GenBank databases. The **Patent** database required more preprocessing, as the heavy and light chains were not stored in the same entry.

Therefore, for each family of patents, only samples with not more than one heavy or heavy and light chains were chosen. If multiple chains appeared, the mapping could produce falsified results, which would decrease the overall quality of the datasets. Due to additional heavy and light chain matching, the quality of the dataset was judged as the “medium” category. The final dataset, **Structures** ^43^, is an aggregate of structure-nanobodies and structures-antibodies, which already includes all necessary information on top of standard structural data. Due to gathering only binding examples without information on binding strength, the boolean type of affinity was chosen, with only “True” values present. However, note that some of the datasets, such as **skempiv2** or **ab-bind** aggregate affinity data from PDB structures, leading to overlaps with this dataset.

The most occurring affinity type is “fuzzy” affinity (stemming from the **buzz** dataset), accounting for 48 % of all entries in ASD. This affinity groups the binding strength into 3 different categories: h (high), m (medium), and l (low). All antibody sequences were required to have complete heavy and light chains, and records missing key identifiers (such as UniProt or PDB IDs for antigens) were either excluded or manually resolved. The intraset deduplication was handled by aggregating rows containing exactly the same antibody and antigens and applying averaging for numerical affinity measurements or mean for categorical expressions. Each source was handled separately in order to ensure the quality of the aggregated dataset.

The averaging was used in datasets such as HIV, met, or COVID-19, where multiple measurements for the same antigen-antibody pairs were present.

After the manual review, each dataset was assigned a confidence label, which represents the quality of the dataset and confidence regarding the source of the dataset. The medium-confidence category includes samples that were included in automatic or semi-automatic ways, including the usage of LLM agents. The high-confidence category comprises well-structured datasets, which require obtaining sequences from external sources such as PDB or UniProt databases ^44,45^. The very high-confidence category represents datasets where affinity, affinity type, and sequences of antigens and antibodies are available without any manual interventions. The curation process resulted in a common structure for all the studies. The antigen datasets contain antibody-antigen pairs along with affinity measurements, structured according to the schema presented in Table 3.

**Table 3:** Description of ASD database schema.

| Column name | Description | Example |
| --- | --- | --- |
| dataset | Dataset abbreviation that allows for reading about its characteristics in this documentation | dlgo |
| heavy_sequence | Heavy chain sequence of the antibody written using amino acids | QVQLVQSGAEVKKPGAS... |
| light_sequence | Light chain sequence of the antibody written using amino acids | DIQMTQSPSTLSASVGD... |
| affinity_type | Type of measurement performed to obtain binding affinity | Ic_50 |
| affinity | numerical value obtained | 0.002 [µg/ml] |
|  | using method described in affinity_type |  |
| antigen_sequence | antigen sequence expressed in amino acid format | MFVFLVLLPLVSSQCVN... |
| metadata.target_uniprot | Id of an antigen connecting it to a specific Uniprot database entry | P0DTC2 |
| metadata.target_pdb | Id of an antigen connecting it to a specific PDB database entry | 7EB0 |
| metadata.target_name | Name or label for the antigen target (e.g., "covid_gamma"). | covid_gamma |
| metadata.source_url | Url pointing to the general resource the sample was obtained from | https://www.pnas.org/doi/10.1073/pnas.2122954119 |
| metadata.heavy_riot_numbering | Outcome of sequence numbering performed using RIOT algorithm on heavy sequence | cdr1_aa='GYTFPTYA',<br>cdr2_aa='INAGNGNT',<br>cdr3_aa='AGGGGRMFQFDYFDY',<br>sequence_alignment_aa='QVQLVQ ... |
| metadata.light_riot_numbering | Outcome of sequence numbering performed using RIOT algorithm on light sequence | cdr1_aa='QSISSW',<br>cdr2_aa='DAS',<br>cdr3_aa='QQYNGYPWT',<br>sequence_alignment_aa='DIQM ... |
| confidence | Level of confidence in binding of exact antibody sequence to an exact antigen sequence | high |
| scfv | Boolean value specifying whether records containing only heavy sequence contain a single-chain variable fragment, or antibodies | False |

## Availability

The dataset is available at https://naturalantibody.com/agab/ for non-commercial applications.

## Results

The curated dataset contains a total of 1,097,946 unique antibody-antigen interaction records, collected from 25 datasets derived from 15 distinct sources. It includes 865,153 unique antibodies and 716,650 complete heavy and light chain pairs. In total, 9,575 unique antigens are represented. These antigens are primarily associated with infectious diseases and cancer. Due to the diverse nature of the database, the number of antibodies associated with each antigen varies from target to target. The most common antigens inside the database are presented in Figure 1.

**Figure 1:**
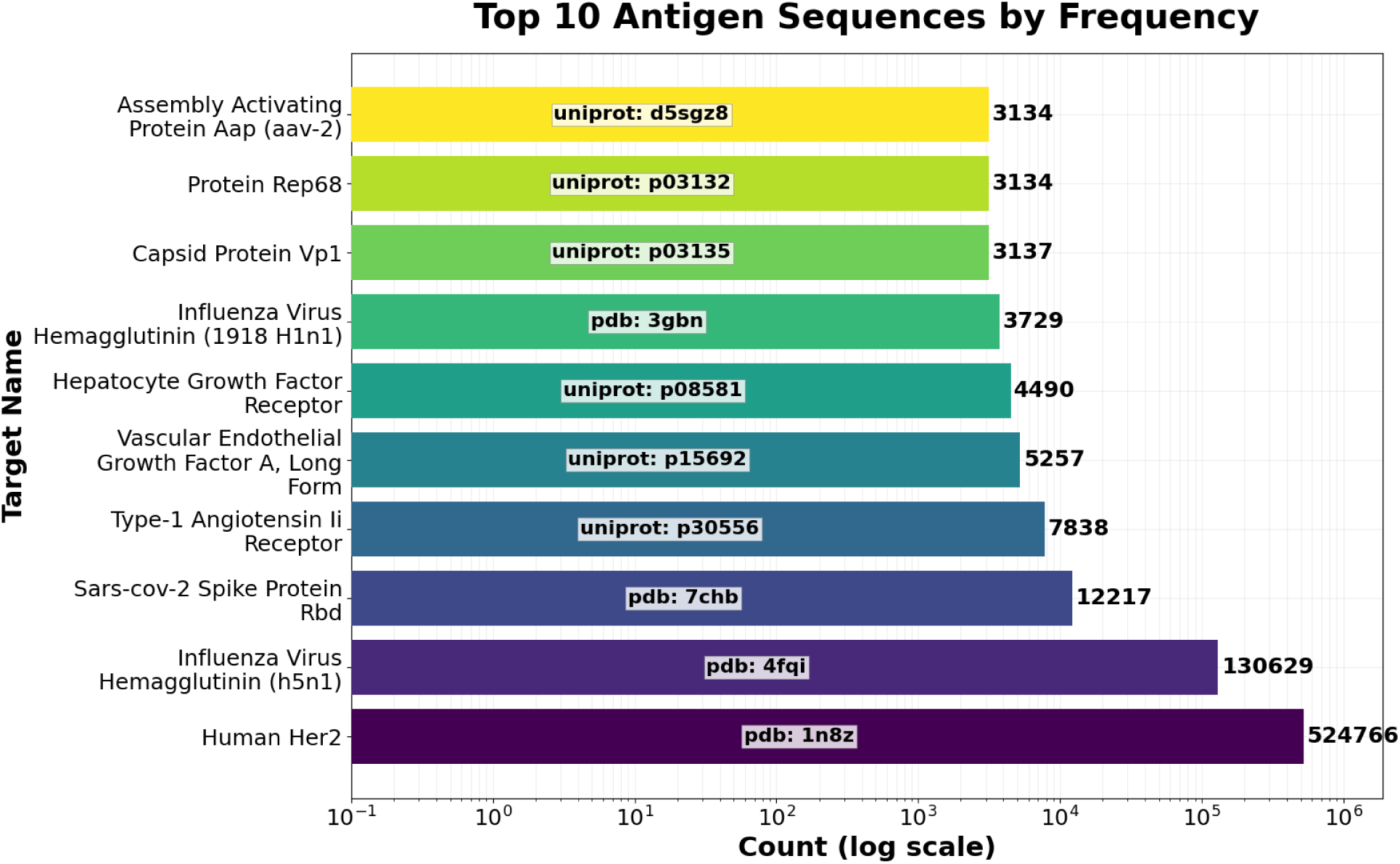
Bar plot with top antigen sequences within the dataset. The counts represent the number of occurrences of each of the selected antigens across all datasets, grouped by specific PDB or UniProt ID to split unique variations of the antigen. The most prevalent antigens are Human Her2 and Influenza Virus.

The uneven distribution of sequence-affinity data across these sub-datasets is the result of heterogeneity of the constituent datasets. For instance, the Buzz collection offers 524,346 entries against a single antigen, making it ideal for ultra-deep mapping of sequence-to-affinity relationships. The Patents dataset, on the other hand, provides 113,117 entries spanning 5,291 unique antigens (≈ approximately 21 per target), supporting broad surveys of cross-reactivity, albeit with limited resolution due to the unreliability of patent data. Other large-scale resources include AbBD (155,853/7, ≈ 22,265 per antigen) and AlphaSeq (198,703/3, ≈ 66,234 per antigen), which combine high depth with narrow antigen scope. Mid-scale datasets offer varying balances of breadth and depth: Cov (27,324/32) delivers ≈ 854 entries per antigen, while HIV (48,008/940) offers ≈ 51 per target. The Literature collection contains 5,636 entries across 884 antigens (≈ 6.4 per target), and GenBank includes 2,989/347 (≈ nine per antigen). Biomap (2,725/594) and SKEMPIv2 (434/21) present broader but shallower landscapes (5 and 21 entries per antigen, respectively). The merged FLAB datasets contribute 6,849 entries across eight distinct antigens (≈ 856 per target), offering focused but high-coverage studies.

Structure-derived resources like structures-antibodies (2,711/1,083) and structures-nanobodies (1,258/390) yield two to three entries per antigen, useful for benchmarking and modeling. Targeted libraries such as AbDesign (672/13, around 52 per target), DLGO (360/10, around 36 per target), AB-Bind (283/10, around 28 per target), and AATP (93/1) address narrower scopes with consistent per-target depth. Several highly focused datasets, such as AAE (35/1), OSH (30/1), MET (4,000/1), and RMNA (10/2), serve niche roles, often centered on single-target studies. The overview of the datasets is given in Figure 2.

**Figure 2:**
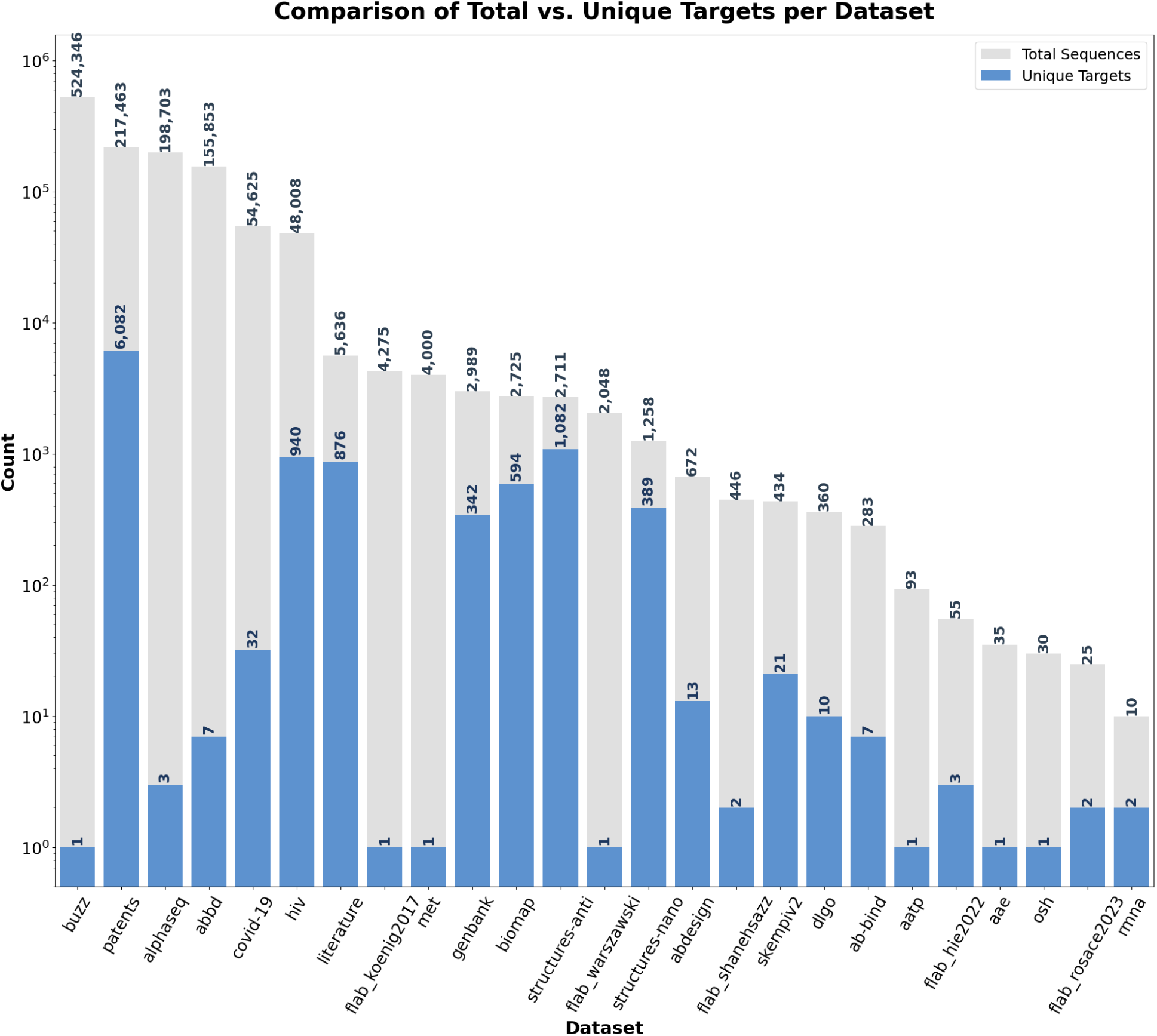
Number of sequences and unique antigens present in each dataset. The blue bars represent the number of unique target antigens in each dataset, while the grey bars account for all interactions within a dataset. The ratio of both bars measures the relative diversity of antigens in the datasets and antibodies in the dataset.

A key limitation of ASD is the substantial imbalance in antigen representation. For instance, the Buzz dataset contributes more than 500,000 entries against a single antigen (HER2), while other antigens are represented by only a handful of interactions. This runs the risk of skewing downstream analyses or machine learning models. To mitigate this, we recommend normalization strategies such as downsampling overrepresented antigens, weighting examples inversely by antigen frequency, or stratified train/test splitting to ensure balanced evaluation. These approaches allow users to tailor ASD to their application, whether prioritizing generalization across antigens or depth within a single target.

A further issue is the overlap between some of our collected datasets. In order to reliably reflect antibody and antigen diversity, we have measured the overlap between existing datasets to allow further pruning of the dataset according to individual needs. The top 10 results are available in Table 4.

**Table 4:**
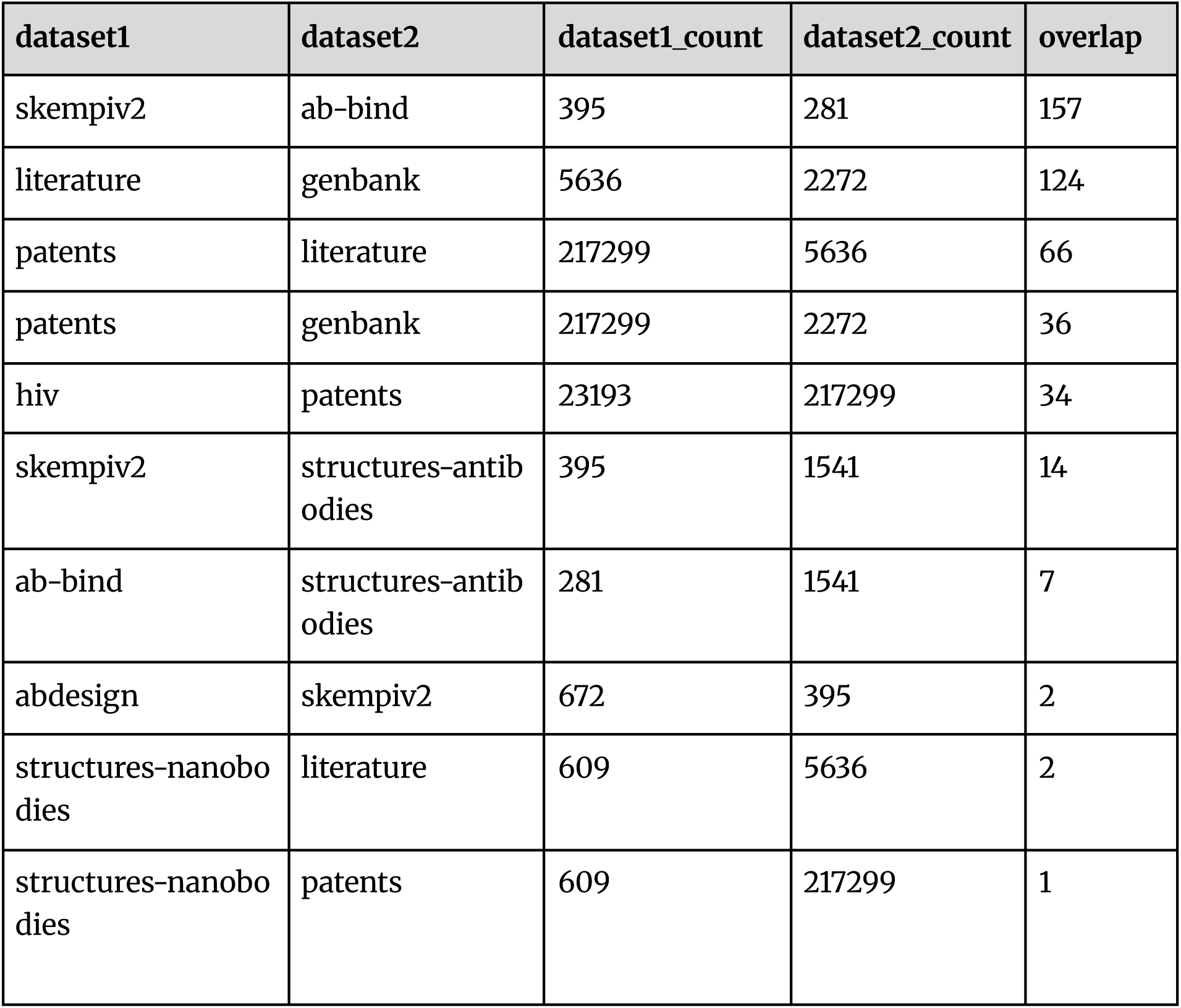
Number of antibody-antigen pairs repeated between datasets.

In addition to antigen diversity, the dataset captures a wide range of ways to capture binding behavior, which we refer to as ‘affinity’ for a lack of better wording. These include quantitative metrics such as Gibbs free energy changes, kinetic constants, and IC₅₀ values, as well as qualitative assessments of binding. Due to variations in experimental methodologies and reporting standards, the frequency of different affinity types may vary. The distribution of the top affinity types across the dataset is shown in Figure 3. Some of the affinity types also include experimental methods (such as ELISA). In cases of some measurements, such as kd or delta_g, the exact methodology for obtaining the data was unknown at the time of writing the paper. The number of sequences in a given category can be distorted by differences in constituent source size. The counts, along with the appropriate datasets, are present in Table 5.

**Figure 3:**
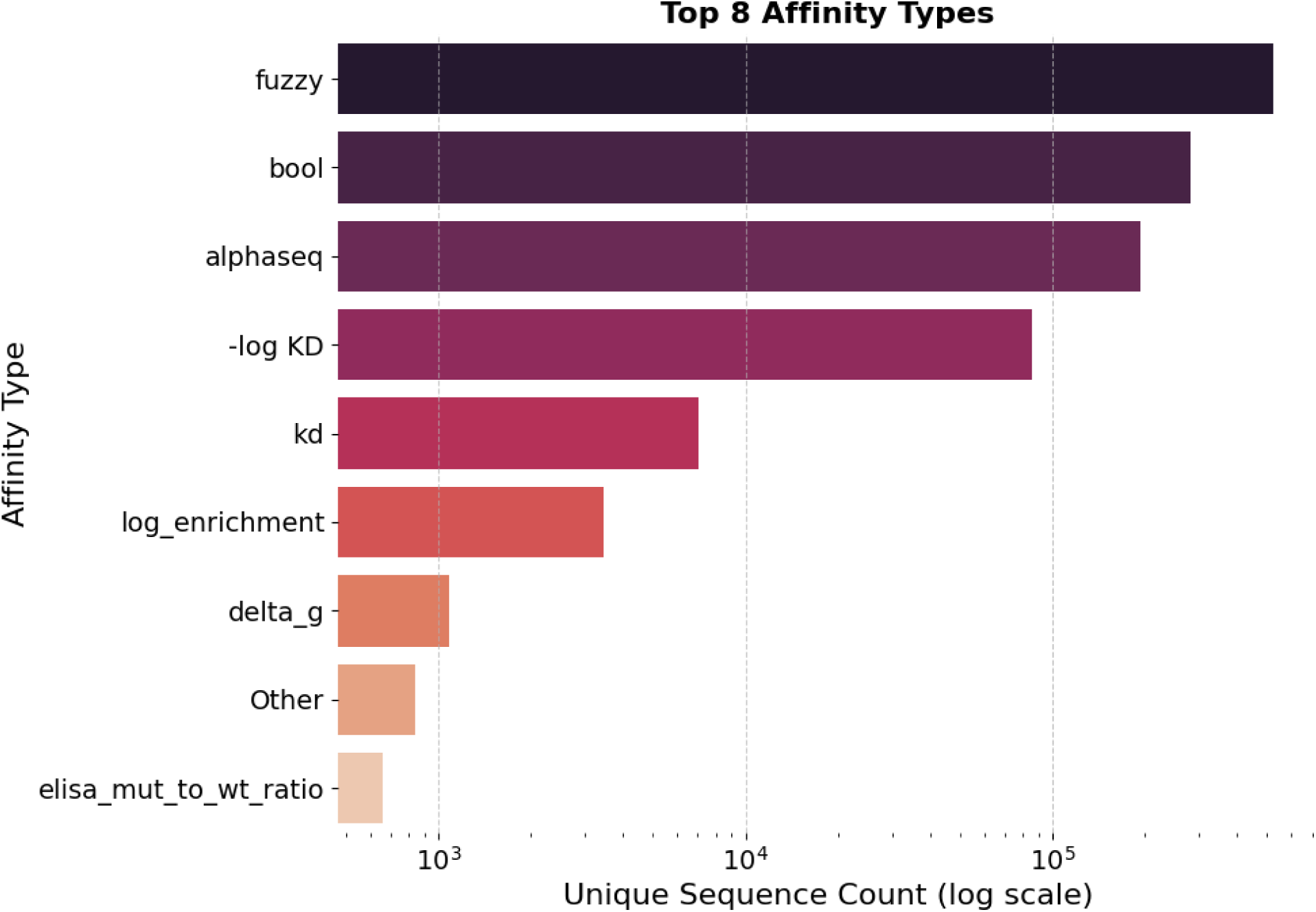
Number of sequences whose binding affinity refers to one of the present categories. The most common category is bool (only informs of binding, not its strength). The affinity that belongs to the most datasets outside of bool is kd, which is often present in smaller datasets of higher quality.

**Table 5:**
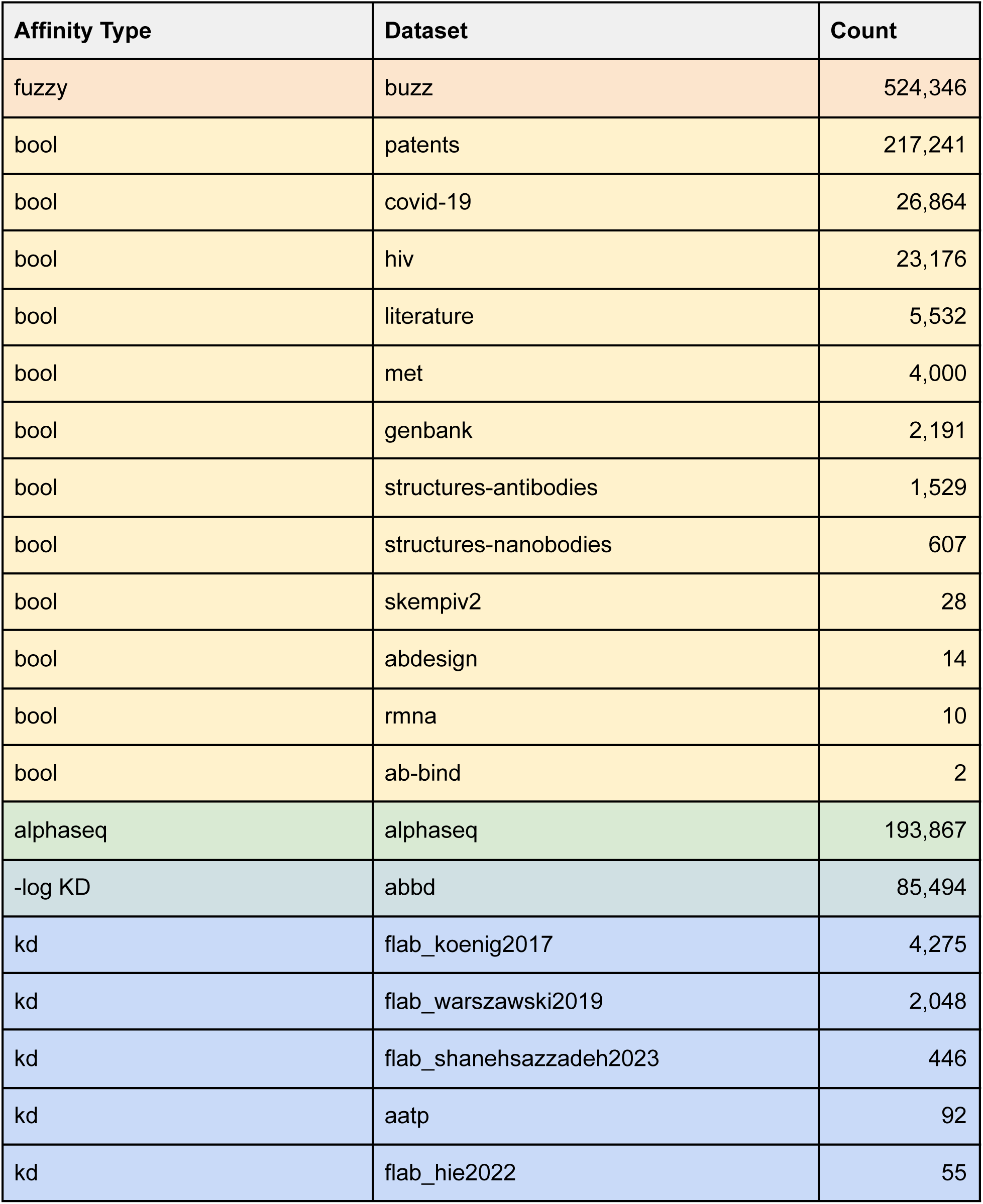
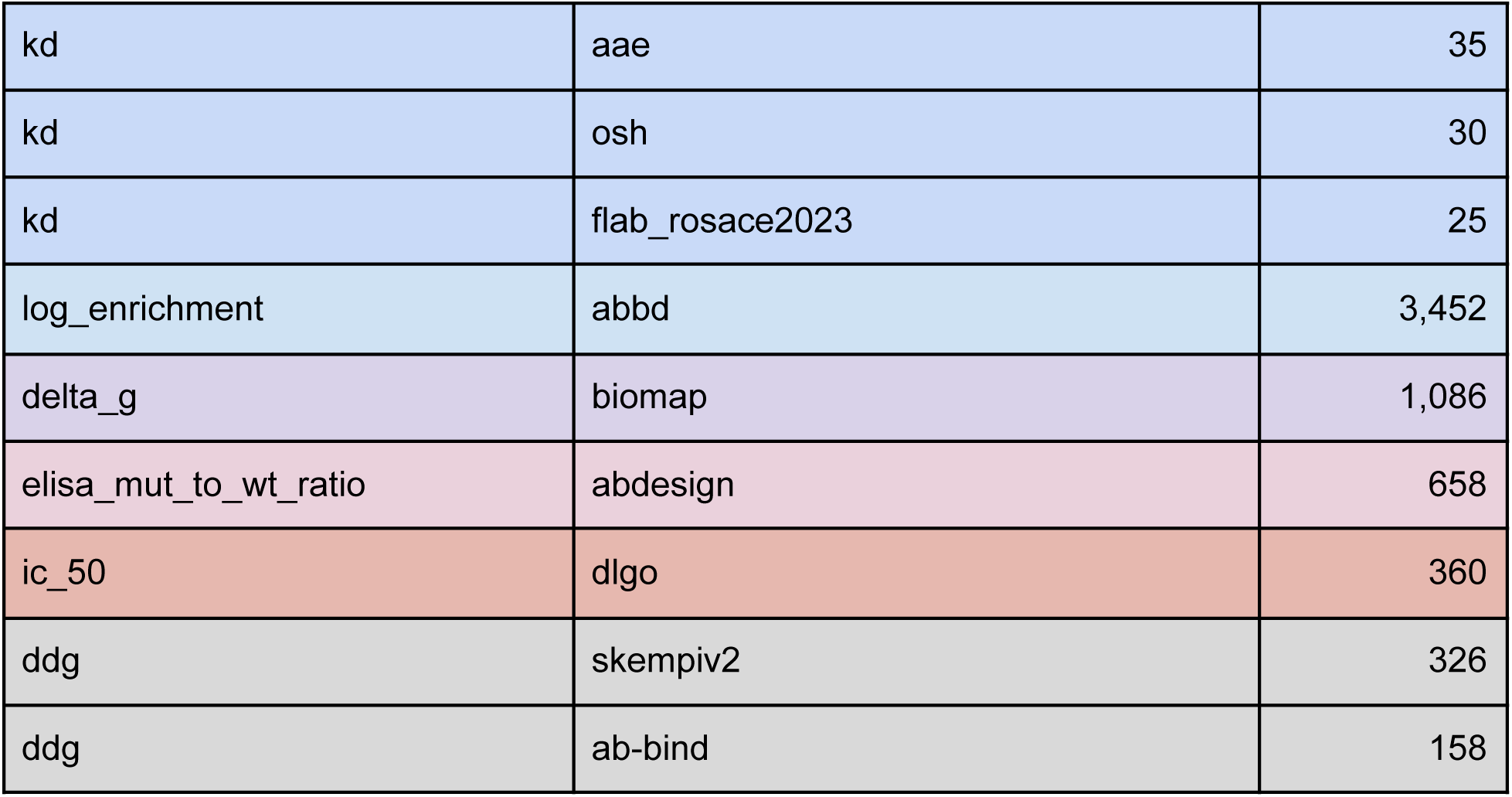
Affinity types differentiated by source dataset, ordered by unique antibody count of the category. The count represents only unique sequences, not all measurements in the dataset, therefore, the numbers can be smaller than the total number of samples in the dataset.

Table 6 summarizes the structural representation of antibody sequences by chain composition. A majority of entries include both heavy and light chains. A smaller subset contains separate heavy or light chains, nanobodies, and SCFV formats. The diversity in sequence types allows for the comparison of both single-domain and standard antibodies. This variation reflects differences in experimental design, sequencing coverage, and the inclusion of non-canonical antibody formats across source datasets.

**Table 6:** Distribution of antibody types based on the chain components. The majority of antibodies are monoclonal antibodies, consisting of both heavy and light chains. A substantial portion contains only heavy chains, representing mostly nanobodies and single-chain variable fragment antibodies, indicating the diversity of the dataset.

| Chain components | Count |
| --- | --- |
| Heavy + Light | 648312 |
| Heavy | 370569 |
| Light | 79065 |

Sequences containing both chains are typically required for full modeling of antibody-antigen interactions, whereas single-chain entries may be suited for focused analyses, such as studies on variable domains or paratope prediction in nanobodies. The observed distribution can also inform prospect preprocessing decisions in downstream pipelines, such as whether to filter for complete paired sequences or to include single-chain entries for broader representation.

The formats are further associated with disparate sequence lengths of antibodies, given in Table 7. Sequence length was computed by summing the number of amino acids in the heavy and light chain sequences for each antibody entry, excluding cases where one or both chains were missing. The resulting values were then binned in different categories based on the distribution of the antibody lengths. The majority of sequences fall into the 124-227 range, which is consistent with variable region only or Fab paired-chain entries typically seen in standard antibody constructs. A smaller fraction of sequences is classified as Short or Medium, reflecting partial records or single-chain formats such as nanobodies. The heterogeneous nature of the database allows for a careful selection of data points based on required characteristics for future research, such as similarity to a tested antibody, developability, or a type of sequence.

**Table 7:** Antibody length distribution by format. Sequences were trimmed to variable regions segments based on germline alignments with RIOT ^46^. The bins are created based on percentile ranges of the combined heavy and light chain sequence length distribution. Nanobodies are predominantly in the shortest length bin (0-115), reflecting their single-domain structure. scFv molecules are present mostly in intermediate lengths (115-227), while monoclonal antibodies and other molecules dominate longer ranges.

| Bucket Size | Nanobodies | scFv | Other |
| --- | --- | --- | --- |
| 0-115 | 5818 | 4306 | 99966 |
| 115-124 | 26973 | 62160 | 77291 |
| 124-227 | 76742 | 61018 | 583347 |
| > 227 | 0 | 262 | 100063 |

Following analysis of the sequence distribution, the germline analysis was performed to reveal the biases of organism assignment in the dataset. Our working hypothesis was that most data in our dataset is human germline-wise. In order to determine V, C, and J calls along with locus species, the RIOT framework was used. Each antibody chain was subjected to separate analysis, resulting in the most probable species assignment. As of the moment of writing this paper, three different species were available for analysis: human, mouse, and alpaca. The majority of antibody chains were assigned to the human germline, followed by smaller contributions from mouse and alpaca (Table 8). The alpaca species was attributed solely to the heavy chain due to a share of nanobodies existing in the dataset.

**Table 8:**
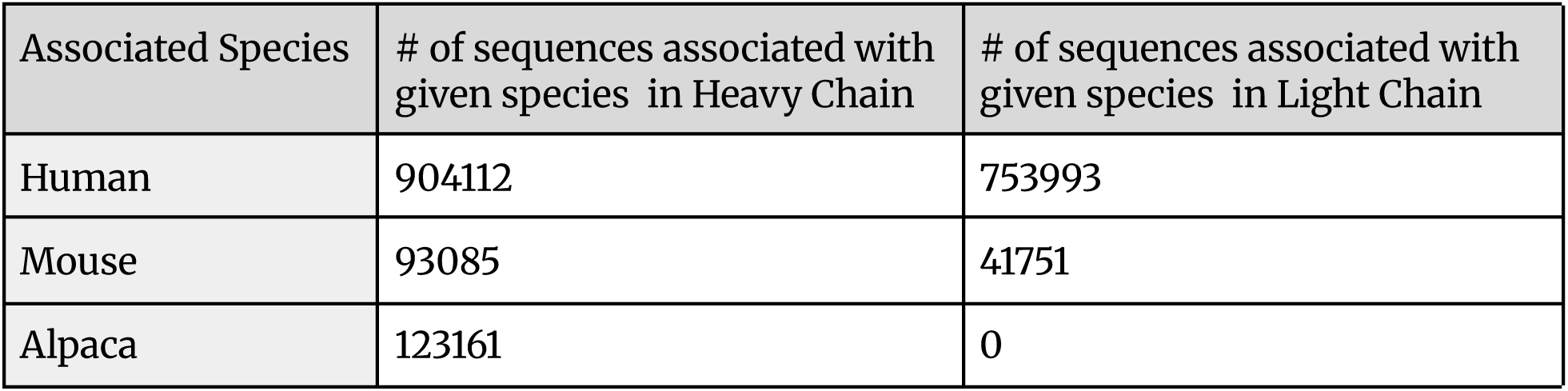
Species distribution of antibody germline assignments. The table shows the number of heavy light chains classified by species using the RIOT framework. Human germlines dominate the dataset, while mouse and alpaca sequences are represented more sparsely.

Following the analysis of germline assignment, a study analysing the most popular genes was performed. The analysis focused on identifying the top five most frequently used V, J, and C genes for both heavy and light chains. For heavy chains, IGHV3-6601 emerged as the dominant V gene, substantially outpacing all other V genes in the dataset. In the J gene analysis, IGHJ402 showed the highest frequency among all heavy chain J genes. The constant region analysis demonstrated IGHE as the predominant C gene for heavy chains. Light chain analysis revealed IGKV1-NL101 as the most abundant V gene, clearly dominating over other light chain V genes. For J genes, IGKJ101 showed the highest usage pattern among light chain J genes. The constant region usage was led by IGKC as the most frequently utilized light chain constant gene. The results shown in Figure 4 clearly show preferences for specific germline genes in both heavy and light chain sequences, with certain genes being used much more frequently than others. The strong selection for particular V, J, and C genes indicates organized usage patterns in the immune system, suggesting that some gene families work better for antibody production, reflected in gene choices for antibody-based therapeutics. However, these observed preferences may be influenced by the types of antigens represented in the dataset, which could affect the apparent gene usage patterns.

**Figure 4:**
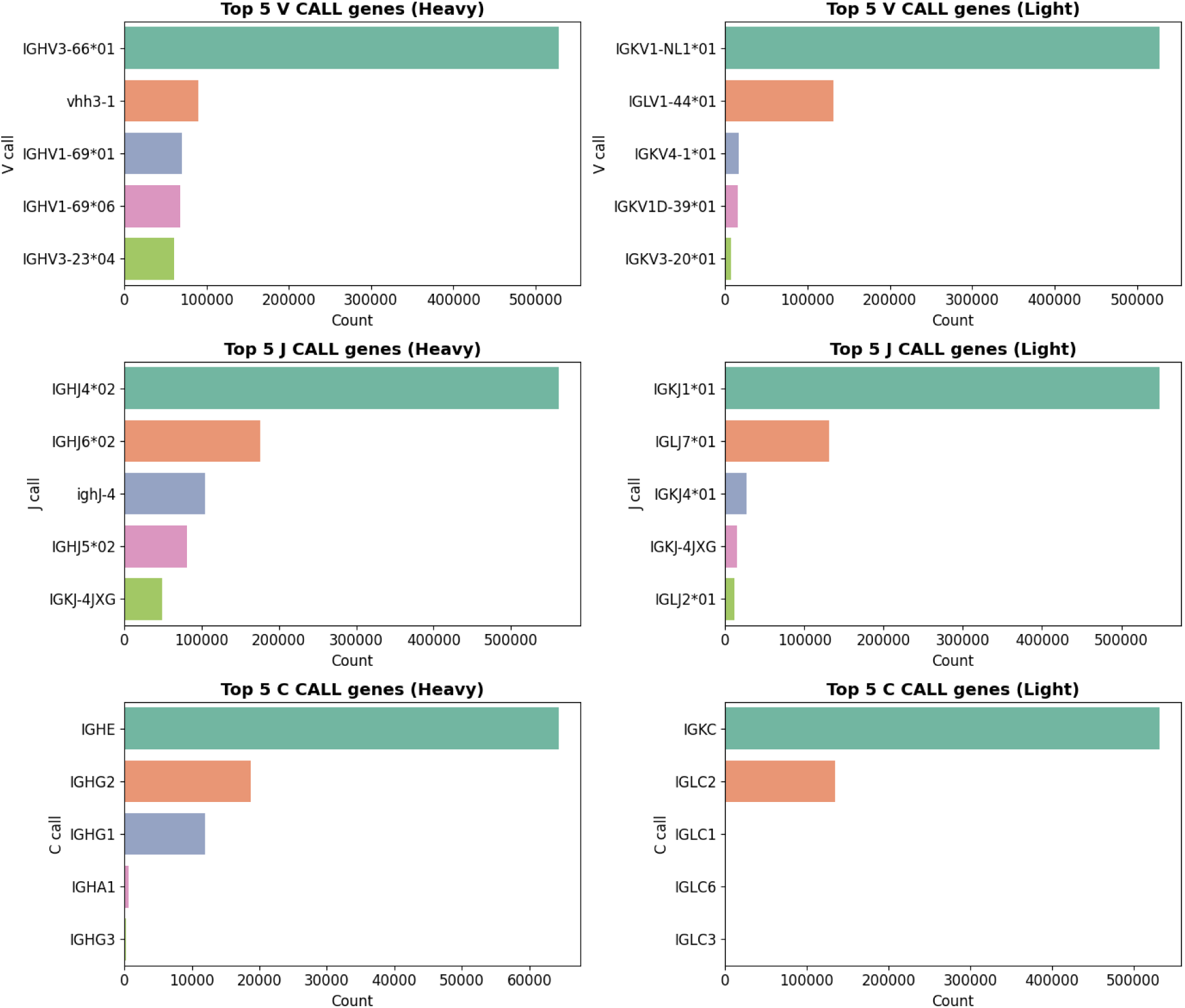
Top germline gene usage across antibody chains. Horizontal bar plots displaying the most frequently utilized V, J, and C genes for heavy and light chains. Each panel shows the top five genes ranked by frequency.

## Discussion

The current standard for obtaining data regarding antibody-antigen modelling consists of performing display or targeted experiments in order to achieve a small number of specific measurements aimed at creating machine learning models specific to a given target. This approach, however, does not allow for generalisation and knowledge transfer between experiments, unless such experiments were performed on a massive scale against a large number of antigens.

Our ASD database addressed both multiple- and single-target development through metadata and diversity in the data sources. It allows both target-specific optimization and generalizable machine learning. The addition of the metadata creates yet another important property of the dataset, data provenance. The explicit data lineage allows for studying the effects of learning within the distribution of the individually generated dataset as well as its generalization towards other datasets.

Although the obtained ‘affinities’ are heterogeneous, they reflect real-world research measurements, tending to group into two different categories - boolean and kd measurements. The metadata and affinity types allow for further filtering and selection of appropriate samples to aid further development. Despite its constraints, ASD provides a flexible and diverse resource that can advance digital drug design. Foremost, it is a reflection of the data that is produced across diverse labs, indicating what benchmarks any modeling solution would have to face.

As a result of its heterogeneity, the key limitation of the dataset is the uneven distribution between different kinds of affinities, antigens, and labels. Most of the Boolean-type affinities include successful bindings, which may cause latent bias in the trained model (notwithstanding that such boolean labels are not necessarily standardized among each other). Therefore, it is important to perform dataset balancing through available means. The more clean, plentiful datasets become available, the sooner the machine learning tools will be able to deliver on their full potential in antibody discovery ^47,48^.

The ASD database addresses the data scarcity issues that hamper machine learning solutions in digitally aided antibody engineering, albeit by consolidation rather than filling critical gaps in volume and quality. Therefore, we propose that the dataset be used primarily as a reference and benchmarking resource. The further development of the database will be conducted as new datasets become available, providing required resources for efficient, cost-effective, and successful development of therapeutics.

## Notes

https://naturalantibody.com/agab/

### Competing Interest Statement

The authors have declared no competing interest.

